## Supplementary Information for "acCRISPR: An activity-correction method for improving the accuracy of CRISPR screens"

Supplementary Figure 1: acCRISPR analysis of Cas12a growth screens in *Yarrowia lipolytica.*

Supplementary Figure 2: Essential gene comparison to *S. cerevisiae* and *S. pombe.*

Supplementary Figure 3: Performance of acCRISPR on the Cas12a screening dataset with predicted sgRNA activities.

Supplementary Figure 4: acCRISPR corrected Tolerance Scores (TS) for 1.5 M NaCl and pH 2.5 tolerance screens.

Supplementary Figure 5: CRISPR-Cas9 and -Cas12a FS distributions on days 2, 4 and 6.

Supplementary Figure 6: Schematic and sequence information of Cas9 and Cas12a amplicons for NGS.

Supplementary Table 1: CS threshold data for Cas9 and Cas12a screens.

Supplementary Table 2: Yeast strains used in this study.

Supplementary Table 3: Plasmids used for genome wide CRISPR screens.

Supplementary Table 4: Sequences of primers used in this study.

Supplementary Table 5: Transformation efficiencies measured as x$10^6$ transformants, for all replicates in the control and treatment strains.

Supplementary Table 6: Primers used for NGS fragment amplification (Cas12a).

Supplementary Table 7: Primers used for NGS fragment amplification (Cas9).

Supplementary Table 8: Parameters for bioinformatics tools used in the analysis of NGS reads (Cas12a).

Supplementary Table 9: Parameters for bioinformatics tools used in the analysis of NGS reads (Cas9).

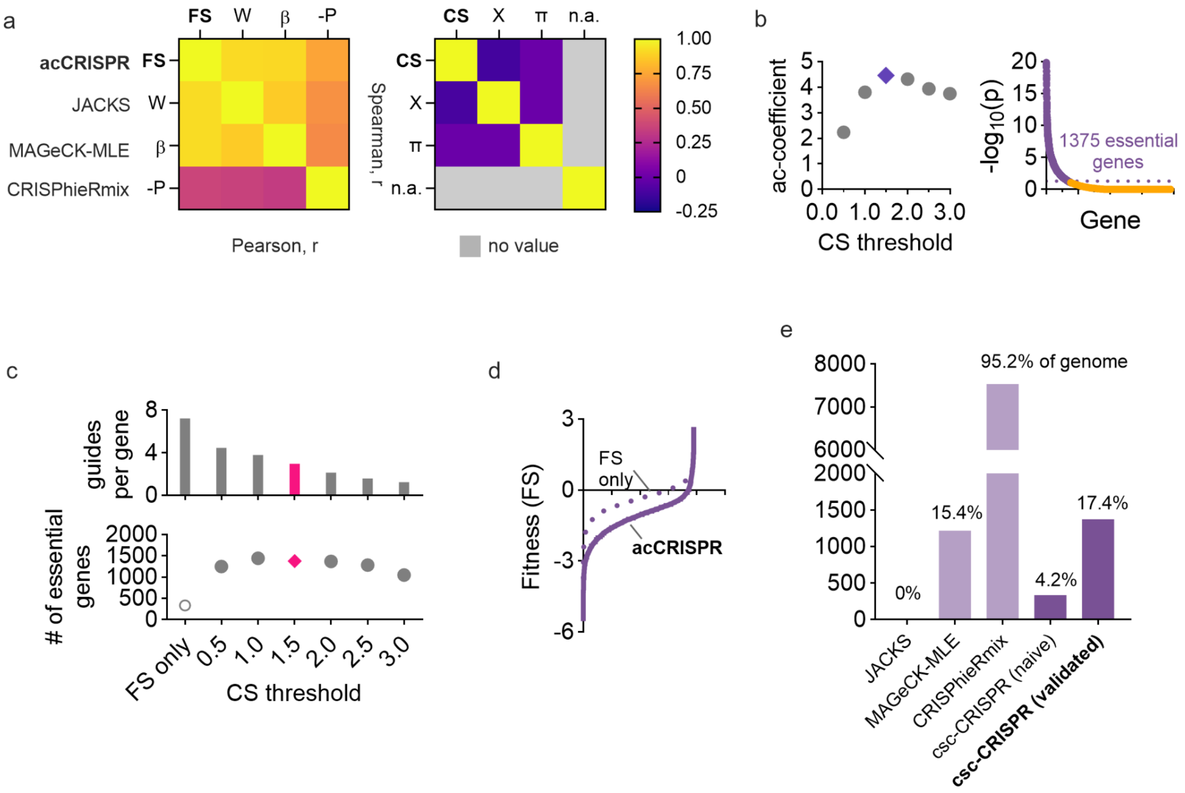


**Supplementary Figure 1. acCRISPR analysis of Cas12a growth screens in *Yarrowia lipolytica*.** (**a**) Heat-maps showing pearson (below diagonal) and spearman (above diagonal) coefficients of fitness effects (FS, W, β & -P; left) and sgRNA cutting efficiencies (CS, X and π; right) and from acCRISPR and three established essential gene identification algorithms, JACKS, MAGeCK-MLE and CRISPhieRmix. **(b)** ac-coefficient is calculated with increasing CS threshold values and maximum value is represented by the purple datapoint. Genes with a p-value < 0.05 were classified as essential at the maximum ac-coefficient value. (**c**) Average number of sgRNA per gene and the number of essential genes predicted with increasing CS threshold. The number of essential genes predicted for the corrected and uncorrected analyses. The data points colored in pink are the guides per gene and number of essential genes determined at the optimum CS threshold. (**d**) Fitness scores of genes with (solid line) and without (dashed line) acCRISPR processing with a CS threshold of 1.5. (**e**) Number of essential genes identified by JACKS [1], MAGeCK-MLE [2], CRISPhieRmix [3], uncorrected FS, and acCRISPR along with the percentage of total genes in the genome are reported.

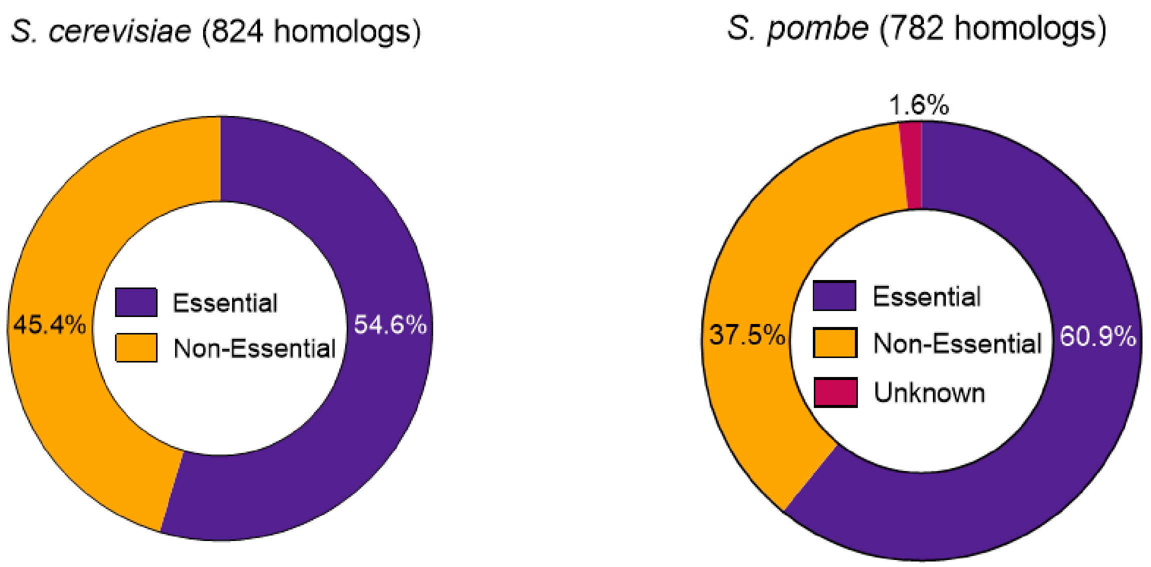


**Supplementary Figure 2. Essential gene comparison to *S. cerevisiae* and *S. pombe*.** Pie charts indicating the percentage of homologs in the *Y. lipolytica* consensus set that are essential, non-essential and have unknown essentiality in *S. cerevisiae* (824 homologs) and *S. pombe* (782 homologs).

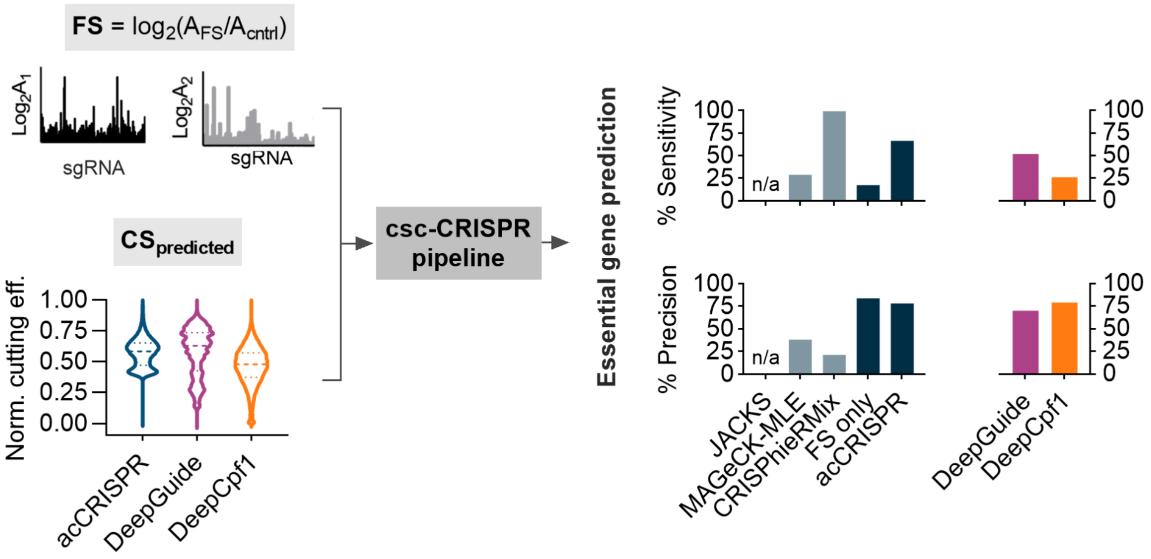


**Supplementary Figure 3. Performance of acCRISPR on the Cas12a screening dataset with predicted sgRNA activities.** Essential genes were determined with acCRISPR utilizing FS along with predicted sgRNA activities from DeepGuide [4] and DeepCpf1 [5]. The violin plot shows min-max normalized sgRNA activity distributions of experimental CS determined by acCRISPR and those from DeepGuide and DeepCpf1. The % sensitivity and % precision in identifying genes from the consensus set is shown (right). Bars indicate the values of these two metrics for each prediction tool as well as for JACKS [1], MAGeCK-MLE [2], CRISPhieRmix [3], uncorrected FS (FS only) and acCRISPR.

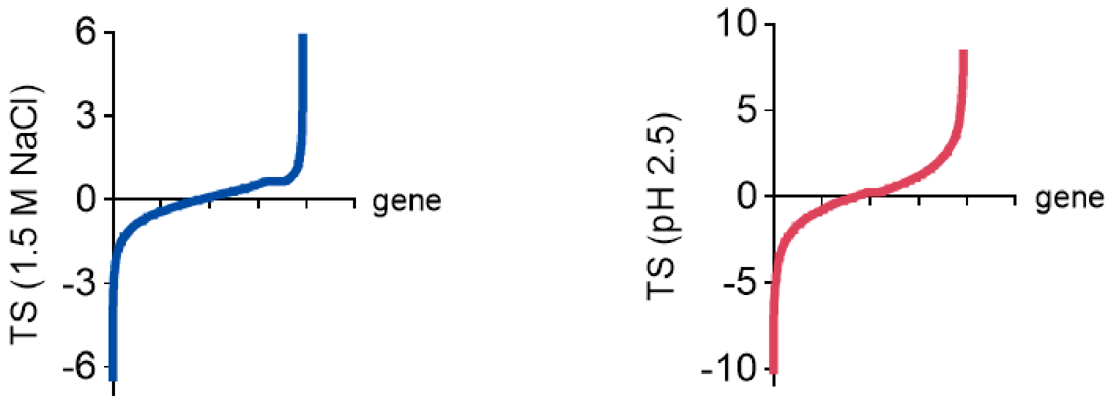


**Supplementary Figure 4. acCRISPR corrected Tolerance Scores (TS) for 1.5 M NaCl and pH 2.5 tolerance screens.** S-curves showing tolerance scores of genes at a CS threshold of 4.5 for two stress conditions - 1.5 M NaCl (left) and pH 2.5 (right).

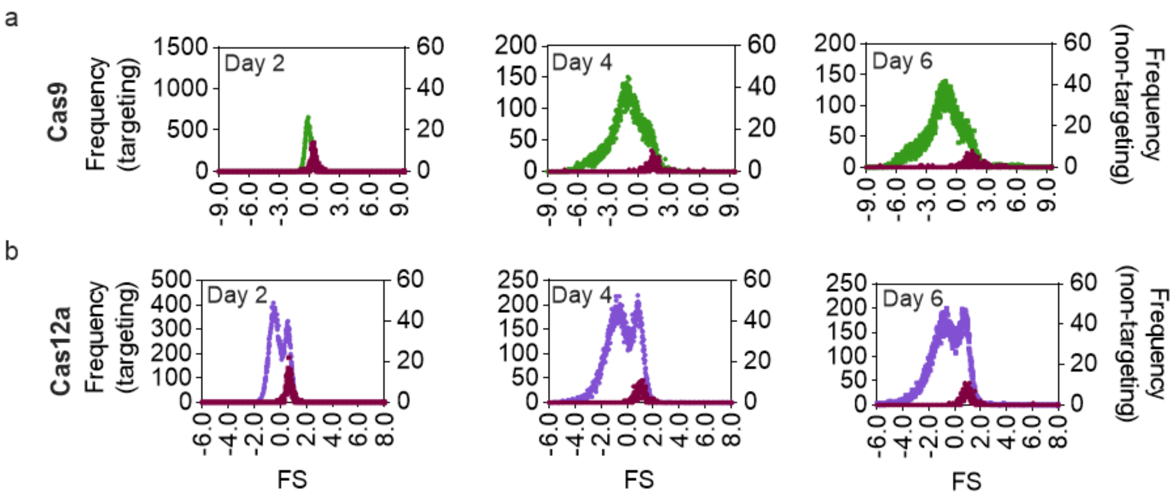


**Supplementary Figure 5. CRISPR-Cas9 and -Cas12a FS distributions on days 2, 4 and 6.** Green and purple distributions plotted on the left y-axis show FS of all targeting sgRNA in the library, while the dark red distributions plotted on the right y-axis represents the non-targeting populations. (a) Histogram of sgRNA FS values in the Cas9 dataset. (b) Histogram of sgRNA FS values in Cas12a dataset.

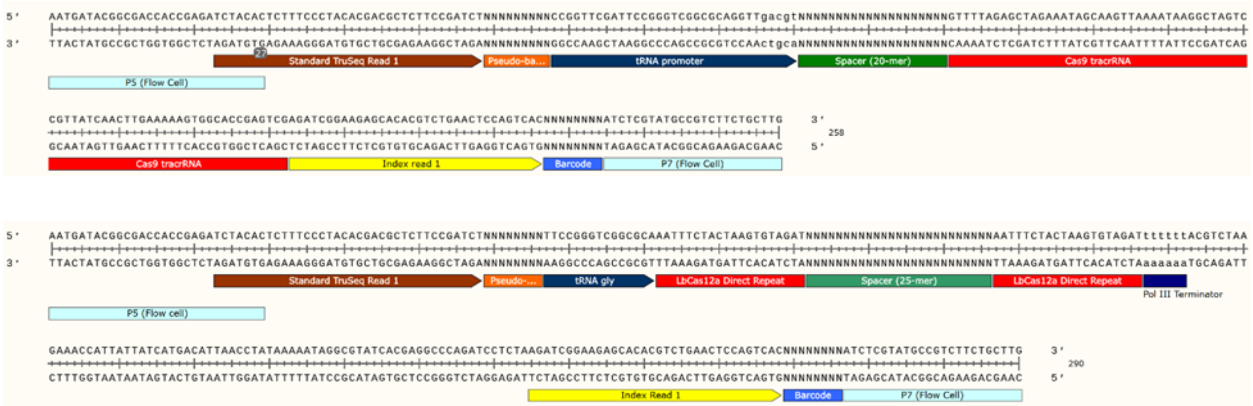


**Supplementary Figure 6. Schematic and sequence information of Cas9 (top) and Cas12a (bottom) amplicons for NGS**. Amplicons contain: (i) P5 and P7 sequences (light blue) that are necessary for binding with the flow cell in Illumina sequencers, (ii) TruSeq adapter (brown) for binding of the sequencing primer, (iii) a portion of $tRNA^{gly}$ (black) expressing the sgRNA, (iv) Cas9 or Cas12 spacer (green) (v) Cas12a associated direct repeats or a portion of the Cas9 tracrRNA sequence (red), (vi) Universal 8 bp Illumina barcodes (blue), (vii) Index read 1 sequence for the binding of primers to sequence the Illumina barcodes, and (viii) 4-9 nt pseudo-barcodes (orange) at the 5' end between the TruSeq and $tRNA^{gly}$ which help demultiplex replicates that contain the same Illumina barcode.

**Supplementary Table 1.** CS threshold data for Cas9 and Cas12a screens. The CS threshold values used to generate 'CS-corrected' libraries and the optimum cutoff value for Cas9 and Cas12a datasets.

| **Cas9 Screen** | **Value** | | | |
|---|---|---|---|---|
| **Cutting efficiency score** | **Lowest cutoff** | **Highest cutoff** | **Step size** | **Optimum cutoff** |
| Experimental CS | 0.5 | 6.0 | 0.5 | 4.5 |
| DeepGuide CS | 0.5 | 6.0 | 0.5 | 4.0 |
| Designer v1 | 0.108 | 0.892 | 0.098 | 0.402 |
| Designer v2 | 20.209 | 78.441 | 7.279 | 49.325 |
| CRISPRspec | 1.215 | 39.175 | 4.745 | 15.45 |
| CRISPRscan | 0.491 | 0.739 | 0.031 | 0.553 |
| SSC | 0.301 | 0.789 | 0.061 | 0.484 |
| uCRISPR | 10.045 | 90.005 | 9.995 | 70.015 |

| **Cas12a Screen** | **Value** | | | |
|---|---|---|---|---|
| **Cutting efficiency score** | **Lowest cutoff** | **Highest cutoff** | **Step size** | **Optimum cutoff** |
| Experimental CS | 0.5 | 3.0 | 0.5 | 1.5 |
| DeepGuide CS | 0.5 | 2.5 | 0.5 | 1.0 |
| DeepCpf1 | 10 | 90 | 10 | 40 |

**Supplementary Table 2.** Yeast strains used in this study.

| Yeast strain genotype | Phenotype |
|---|---|
| PO1f (MatA, *leu2-270*, *ura3-302*, *xpr2-322*, *axp-2*) | Wild type strain |
| PO1f *Δku70* | PO1f with disrupted KU70, which facilitates the non-homologous end joining DNA repair pathway |
| PO1f UAS1B8-TEF(136)-Cas9 -CycT::A08 | PO1f expressing *Y. lipolytica* codon optimized Cas9 gene at the A08 locus |
| PO1f UAS1B8-TEF(136)-LbCas12a -CycT::A08 | PO1f expressing *Y. lipolytica* codon optimized LbCas12a gene at the A08 locus |
| PO1f *Δku70* UAS1B8-TEF(136)-Cas9 -CycT::A08 | *KU70* disrupted in Cas9 integrated PO1f strain |
| PO1f *Δku70* UAS1B8-TEF(136)-LbCas12a -CycT::A08 | *KU70* disrupted in LbCas12a integrated PO1f strain |

**Supplementary Table 3.** Plasmids used for genome wide CRISPR screens.

| Plasmid name | Reference | Function |
|---|---|---|
| pCpf1_yl | 6 | Plasmid for CRISPR-LbCas12a based gene editing in *Y. lipolytica* |
| pCRISPRyl (Addgene #70007) | 7 | Plasmid for CRISPR-Cas9 based gene editing in *Y. lipolytica* |
| pLbCas12ayl | This study and 4 | Plasmid for CRISPR-LbCas12a based gene editing in *Y. lipolytic*a. sgRNA is flanked on either end by the direct repeat, to allow sgRNAs to end in T residues without being construed as part of the PolyT terminator |
| pHR_A08_hrGFP (Addgene #84615) | 8 | Plasmid containing homology arms for integration of hrGFP into the A08 locus |
| pHR_A08_LbCas12a | This study and 4 | Plasmid containing homology arms for integration of LbCas12a into the A08 locus |
| pHR_A08_Cas9 | 9 | Plasmid containing homology arms for integration of Cas9 into the A08 locus |
| pLbCas12ayl-GW | This study and 4 | Vector containing sgRNA expression cassette for cloning Cas12a sgRNA library. (Does not contain Cas12a expression cassette) |
| pCas9yl-GW | 9 | Vector containing sgRNA expression cassette for cloning Cas9 sgRNA library. (Does not contain Cas9 expression cassette) |
| pCRISPRyl_KU70 | This study and 10 | CRISPR plasmid for the disruption of KU70 |

**Supplementary Table 4.** Sequences of primers used in this study.

| Primer name | Primer Sequence |
|---|---|
| ExtraDR-F | CGGCGCAAATTTCTACTAAGTGTAGACTAGTAATTTCTACTAAGTGTAGATTTTTTTACGTCTAAGAAACCATTATT |
| ExtraDR-R | AATAATGGTTTCTTAGACGTAAAAAAATCTACACTTAGTAGAAATTACTAGTCTACACTTAGTAGAAATTTGCGCCG |
| Cpf1-Int-F | TGCCTGGAGCCGAGTACGGCATTGATTACTAGTCCGGGTTCGAAGGTACCAAG |
| Cpf1-Int-R | TTAGGCTGGGTCTCGAGAGCAAAGAAGCCTAGGGCAAATTAAAGCCTTCGAGCG |
| BRIDGE-F | CTAAATTTGATGAAAGGGGGATCCCCCGGGTGGCGTAATCATGGTCATAGCTGTTTCCTG |
| BRIDGE-R | CAGGAAACAGCTATGACCATGATTACGCCACCCGGGGGATCCCCCTTTCATCAAATTTAG |
| A08-Seq-F | AGCCGAGTACGGCATTGAT |
| A08-Seq-R | TCAATGTAGCCTCCTCCAACC |
| Tef_Seq-F | GTTGGGACTTTAGCCAAG |
| Lb1-R | CTTCTGCTTGGTCTTCTGGTTG |
| Lb2-F | AACCTGTACAACCAGAAGACCAAG |
| Lb3-F | AAGGAGACCAACCGAGACGAG |
| Lb4-F | AACCTGCACACCATGTACTTCAAG |
| Lb5-F | CCAGATCACCAACAAGTTCGAGTC |
| M13-F | GTAAAACGACGGCCAGT |
| InversePCR-F | TTTTTTTACGTCTAAGAAACCATTATTATCATGACATTAACCT |
| InversePCR-R | TGCGCCGACCCGGAATCGAACCGGGGGCCC |
| OLS-F | GTTTAGTGGTAAAATCCATCGTTGCCATCG |
| OLS-R | GATACGCCTATTTTTATAGGTTAATGTCATG |
| qPCR-GW-F | TTATGAACTGAAAGTTGATGGC |
| qPCR-GW-R | TCACACAGGAAACAGCTATG |
| Cr_1250 | TATAAGAATCATTCAAAGGCGCGCATGGATAAGAAATACTCCATTGGCCTG |
| Cr_1254 | ATAACTAATTACATGAGGCTAGCTTACAGCATGTCCAGATCGAAATCG |

**Supplementary Table 5.** Transformation efficiencies measured as x$10^6$ transformants, for all replicates in the control and treatment strains.

| **Cas9 Screen** | **Replicate Transformation Efficiency (x$10^6$ transformants)** | | |
|---|---|---|---|
| **Strain** | **R1** | **R2** | **R3** |
| PO1f | 12.35 | 11.39 | 15.80 |
| PO1f Cas12a | 11.42 | 8.29 | 10.64 |
| PO1f Cas12a Δku70 | 6.79 | 7.33 | 7.08 |

| **Cas12a Screen** | **Replicate Transformation Efficiency (x$10^6$ transformants)** | | |
|---|---|---|---|
| **Strain** | **R1** | **R2** | **R3** |
| PO1f Δku70 | 6.89 | 6.21 | 5.43 |
| PO1f Cas12a Δku70 | 5.06 | 4.29 | 4.41 |
| PO1f | 11.93 | 8.28 | 4.23 |
| PO1f Cas12a | 6.32 | 5.47 | 6.11 |

**Supplementary Table 6.** Primers used for NGS fragment amplification (Cas12a)

| Primer name | Primer Sequence | Illumina Barcode (Reverse primer) / Pseudo-Barcode (Forward primer) for demultiplexing |
|---|---|---|
| ILU1-F | AATGATACGGCGACCACCGAGATCTACACTCTTTCCCTACACGACGCTCTTCCGATCTTTCCGGGTCGGCGCAAATTTC | ^TTCCGG |
| ILU2-F | AATGATACGGCGACCACCGAGATCTACACTCTTTCCCTACACGACGCTCTTCCGATCTAGATCGGGTCGGCGCAAATTTCT | ^AGATCG |
| ILU3-F | AATGATACGGCGACCACCGAGATCTACACTCTTTCCCTACACGACGCTCTTCCGATCTGCTATTCGGGTCGGCGCAAATTTCT | ^GCTATT |
| ILU4-F | AATGATACGGCGACCACCGAGATCTACACTCTTTCCCTACACGACGCTCTTCCGATCTCAGGACTACGGGTCGGCGCAAATTTCT | ^CAGGAC |
| ILU1-R | CAAGCAGAAGACGGCATACGAGATTCGCCTTGGTGACTGGAGTTCAGACGTGTGCTCTTCCGATCTTAGAGGATCTGGGCCTCGTGATAC | CAAGGCGA |
| ILU2-R | CAAGCAGAAGACGGCATACGAGATGACGAGAGGTGACTGGAGTTCAGACGTGTGCTCTTCCGATCTTAGAGGATCTGGGCCTCGTGATAC | CTCTCGTC |
| ILU3-R | CAAGCAGAAGACGGCATACGAGATAGACTTGGGTGACTGGAGTTCAGACGTGTGCTCTTCCGATCTTAGAGGATCTGGGCCTCGTGATAC | CCAAGTCT |
| ILU4-R | CAAGCAGAAGACGGCATACGAGATCTGTATTAGTGACTGGAGTTCAGACGTGTGCTCTTCCGATCTTAGAGGATCTGGGCCTCGTGATAC | TAATACAG |
| ILU5-R | CAAGCAGAAGACGGCATACGAGATCCTGAACCGTGACTGGAGTTCAGACGTGTGCTCTTCCGATCTTAGAGGATCTGGGCCTCGTGATAC | GGTTCAGG |
| ILU6-R | CAAGCAGAAGACGGCATACGAGATATCAGGTTGTGACTGGAGTTCAGACGTGTGCTCTTCCGATCTTAGAGGATCTGGGCCTCGTGATAC | AACCTGAT |
| ILU7-R | CAAGCAGAAGACGGCATACGAGATTAGGTGACGTGACTGGAGTTCAGACGTGTGCTCTTCCGATCTTAGAGGATCTGGGCCTCGTGATAC | GTCACCTA |

| | | |
|---|---|---|
| ILU8-R | CAAGCAGAAGACGGCATACGAGATCGAACAGTGTGACTGGAGTTCAGACGTGTGCTCTTCCGATCTTAGAGGATCTGGGCCTCGTGATAC | ACTGTTCG |
| ILU9-R | CAAGCAGAAGACGGCATACGAGATGTTCGATCGTGACTGGAGTTCAGACGTGTGCTCTTCCGATCTTAGAGGATCTGGGCCTCGTGATAC | GATCGAAC |
| ILU10-R | CAAGCAGAAGACGGCATACGAGATACCTAGCTGTGACTGGAGTTCAGACGTGTGCCTTCCGATCTTAGAGGATCTGGGCCTCGTGATAC | AGCTAGGT |
| ILU11-R | CAAGCAGAAGACGGCATACGAGATAGAGATGAGTGACTGGAGTTCAGACGTGTGCTCTTCCGATCTTAGAGGATCTGGGCCTCGTGATAC | TCATCTCT |
| ILU12-R | CAAGCAGAAGACGGCATACGAGATCTGGACTTGTGACTGGAGTTCAGACGTGTGCTCTTCCGATCTTAGAGGATCTGGGCCTCGTGATAC | AAGTCCAG |

**Supplementary Table 7.** Primers used for NGS fragment amplification (Cas9)

| Primer name | Primer Sequence | Illumina Barcode (Reverse primer) / Pseudo-Barcode (Forward primer) for demultiplexing |
|---|---|---|
| **Cr_1665** | AATGATACGGCGACCACCGAGATCTACACTCTTTCCCTACACGACGCTCTTCCGATCTAGTCCGGTTCGATTCCGGGTC | ^AGTCCG |
| **Cr_1666** | AATGATACGGCGACCACCGAGATCTACACTCTTTCCCTACACGACGCTCTTCCGATCTGTAGTCCGGTTCGATTCCGGGTC | ^GTAGTC |
| **Cr_1667** | AATGATACGGCGACCACCGAGATCTACACTCTTTCCCTACACGACGCTCTTCCGATCTCAGTAGTCCGGTTCGATTCCGGGTC | ^CAGTAG |
| **Cr_1668** | AATGATACGGCGACCACCGAGATCTACACTCTTTCCCTACACGACGCTCTTCCGATCTTCCAGTAGTCCGGTTCGATTCCGGGTC | ^TCCAGT |
| **Cr_1669** | CAAGCAGAAGACGGCATACGAGATTCGCCTTGGTGACTGGAGTTCAGACGTGTGCTCTTCCGATCTCGACTCGGTGCCACTTTTTCAAG | CAAGGCGA |
| **Cr_1670** | CAAGCAGAAGACGGCATACGAGATATAGCGTCGTGACTGGAGTTCAGACGTGTGCTCTTCCGATCTCGACTCGGTGCCACTTTTTCAAG | GACGCTAT |
| **Cr_1671** | CAAGCAGAAGACGGCATACGAGATGAAGAAGTGTGACTGGAGTTCAGACGTGTGCTCTTCCGATCTCGACTCGGTGCCACTTTTTCAAG | ACTTCTTC |
| **Cr_1672** | CAAGCAGAAGACGGCATACGAGATATTCTAGGGTGACTGGAGTTCAGACGTGTGCTCTTCCGATCTCGACTCGGTGCCACTTTTTCAAG | CCTAGAAT |
| **Cr_1673** | CAAGCAGAAGACGGCATACGAGATCGTTACCAGTGACTGGAGTTCAGACGTGTGCTCTTCCGATCTCGACTCGGTGCCACTTTTTCAAG | TGGTAACG |
| **Cr_1709** | CAAGCAGAAGACGGCATACGAGATGTCTGATGGTGACTGGAGTTCAGACGTGTGCTCTTCCGATCTCGACTCGGTGCCACTTTTTCAAG | CATCAGAC |
| **Cr_1710** | CAAGCAGAAGACGGCATACGAGATTTACGCACGTGACTGGAGTTCAGACGTGTGCTCTTCCGATCTCGACTCGGTGCCACTTTTTCAAG | GTGCGTAA |

**Cr_1711** CAAGCAGAAGACGGCATACGAGATTTGA CTATTCAA
ATAGGTGACTGGAGTTCAGACGTGTGCTC
TTCCGATCTCGACTCGGTGCCACTTTTTC
AAG

**Supplementary Table 8.** Parameters for bioinformatics tools on Galaxy [11] used in the analysis of NGS reads (Cas12a)

| Tool | Version | Parameters* |
|---|---|---|
| FastQC | v0.11.8 | Default settings |
| Cutadapt | Galaxy Version 1.16.6 [12] | The 3 biological replicates of a given sample at a given time-point in the Cas12a screen always had the same reverse primer containing the Illumina barcode, and forward primers ILU1-F, ILU3-F and ILU4-F; or ILU2-F, ILU3-F and ILU4-F each containing different pseudo-barcodes. Thus Cutadapt was used to demultiplex biological replicates from each other.<br>▪ 5' (Front) anchored 6 bp pseudo-barcodes to be demultiplexed (-g): ^NNNNNN (refer to previous table for pseudo-barcode-forward primer association).<br>▪ Maximum error rate (--error-rate): 0.2<br>▪ Match times (--times): 1<br>▪ Minimum overlap length (--overlap): 4<br>▪ Multiple output: Yes (Each demultiplexed readset is written to a separate file) |
| Trimmomatic | v0.38 | ▪ HEADCROP: 29 (if amplified by ILU1-F); or 30 (if amplified by ILU2-F); or 32 (if amplified by ILU3-F); or 34 (if amplified by ILU4-F)<br>▪ CROP: 25 |
| Bowtie2** | v2.4.2 | ▪ Number of allowed mismatches in seed alignment (-N): 1<br>▪ Length of the seed substring (-L): 21<br>▪ Function governing interval between seed substrings in multiseed alignment (-i): S,1,0.50<br>▪ Function governing maximum number of ambiguous characters (--n-ceil): L,0,0.15<br>▪ Alignment mode: end-to-end<br>▪ Number of attempts of consecutive seed extension events (-D): 20<br>▪ Number of times re-seeding occurs for repetitive reads: 3<br>▪ Save mapping statistics: Yes |

* All parameters other than those mentioned here are kept at default values.
** Bowtie2 usage needs a genome fasta file for alignment. Nontargeting sgRNA and any other sgRNA that Bowtie2 could not find within the original CLIB89 genome file were appended as an extra chromosome so that Bowtie could align all sgRNA for the purposes of generating counts.

**Supplementary Table 9.** Parameters for bioinformatics tools on Galaxy [11] used in the analysis of NGS reads (Cas9)

| Tool | Version | Parameters* |
|---|---|---|
| FastQC | v0.11.8 | Default settings |
| Cutadapt | Galaxy Version 1.16.6 [12] | Cutadapt was used to demultiplex samples containing the same Illumina barcode, but different pseudobarcodes at the 5' end of the read. Samples were amplified with reverse primers Cr1669-1673;Cr1709-1711 and forward primers Cr1665-1668 each containing a different pseudo barcode as mentioned in Table<br>▪ 5' (Front) anchored 6 bp pseudo-barcodes to be demultiplexed (-g): ^NNNNNN (refer to previous table for pseudo-barcode-forward primer association).<br>▪ Maximum error rate (--error-rate): 0.2<br>▪ Match times (--times): 1<br>▪ Minimum overlap length (--overlap): 4<br>▪ Multiple output: Yes (Each demultiplexed readset is written to a separate file) |
| Trimmomatic | v0.38 | ▪ HEADCROP: 30 (if amplified by Cr1665); or 32 (if amplified by Cr1666); or 34 (if amplified by Cr1667); or 36 (if amplified by Cr1668)<br>▪ CROP: 20 |
| Bowtie2** | v2.4.2 | ▪ Number of allowed mismatches in seed alignment (-N): 1<br>▪ Length of the seed substring (-L): 19<br>▪ Function governing interval between seed substrings in multiseed alignment (-i): S,1,0.50<br>▪ Function governing maximum number of ambiguous characters (--n-ceil): L,0,0.15<br>▪ Alignment mode: end-to-end<br>▪ Number of attempts of consecutive seed extension events (-D): 20<br>▪ Number of times re-seeding occurs for repetitive reads: 3<br>▪ Save mapping statistics: Yes |

* All parameters other than those mentioned here are kept at default values.
** Bowtie2 usage needs a genome fasta file for alignment. Nontargeting sgRNA and any other sgRNA that Bowtie2 could not find within the original CLIB89 genome file were appended as an extra chromosome so that Bowtie could align all sgRNA for the purposes of generating counts.
